# Coexistence of Virome-Encoded Human Probiotic Genes and Pathogenic Genes in Global Habitats

**DOI:** 10.1101/2025.01.10.632382

**Authors:** Min Qian, Dong Zhu, Ke-yu Yao, Shu-yue Liu, Meng-ke Li, Mao Ye, Yong-guan Zhu

**Author notes:** **Corresponding autho**r: Mao Ye.

## Abstract

Some probiotic genes for humans obtained through horizontal gene transfer (HGT) have been screened in viral genomes, yet the occurrence mode, geographic distribution, and functional characteristics of these genes in viruses and the coexistence between pathogenic virulence genes and probiotic genes in the viral genome, remain insufficiently understood and verified. In this study, using IMG/VR v4 database, 4,556 viruses carrying probiotic gene segments have been discovered in eight habitat types (humans, animals, freshwater, marine, other water bodies, soil, and plants), 13 regions of 76 countries around the world. Among them, the 478 viruses carrying probiotic genes that distributed in humans account for the highest proportion among the viruses with predictable hosts. In viruses carrying probiotic genes, *BCO1*, *bioB*, *COQ2*, *GPX1*, *GSTs*, *GSTT1*, *GULO,* and *menA* are not only probiotic genes encoded by viruses, but also auxiliary metabolic genes. Four virulence genes harmful to humans in viral sequences carrying probiotic genes with transcription and expression potential, have been found, indicating that virulence genes encoded by viruses can coexist with probiotic genes. Further metagenome sequencing revealed the consistency between the probiotic gene asd in the bacillus prophage and the corresponding phage auxiliary metabolic genes. This study integrates probiotic gene databases into viral genomics, revealing the compatibility between viral probiotic and virulence genes, and offering new insights for viral applications.

**IMPORTANCE:** Probiotic genes acquired by humans through horizontal gene transfer (HGT) have been identified within viral genomes. However, their occurrence, distribution, and functions, along with the coexistence of pathogenic and probiotic genes in viruses, remain poorly understood. This study advocates for the utilization of probiotic gene databases in viral genomes and innovatively elucidates the phenomenon of compatibility between viral probiotic genes and virulence genes, offering fresh perspectives on the diverse applications of viruses. By promoting the application of these databases, the study reveals the compatibility between probiotic and pathogenic genes in viruses, providing novel and insightful implications for the various utilizations of viruses.

## Main

The relationship between viruses and humans is complex, spanning from beneficial to antagonistic(1–3). Viruses have fought against humans for over 300,000 years on several levels(2), from host adaptation to evolution(4) and carrier propagation to infection loading(5). Throughout human history, virus-induced diseases have caused several global public health crises. Some events, from the plague in Europe in the 14^th^ century(6) to Spanish influenza in the 20^th^ century(7) and HIV/AIDS in the 21^st^ century(8) and then to the recent COVID-19 epidemic(9), exhibited the pathogenicity of viruses and their risks to human health. The pathogenicity of viruses is mainly due to virulence genes in the genome(10, 11). Virulence genes enhance the pathogenicity of viruses through multiple mechanisms, destroy the structures and functions of host cells, and may cause several diseases(10–13).

Nevertheless, some genes in favor of humans have been developed in the viral genome through the competition between viruses and humans over mellennia(14, 15). Some studies have demonstrated that some viruses carry virulence genes and contain probiotic gene segments(15–17). These probiotic genes are integrated from host cells into the viral genome through horizontal gene transfer (HGT), forming auxiliary metabolic genes (AMGs)(18–21). These AMGs not only help viruses adapt to different environments but can also positively affect hosts during infection(21–24). This discovery provides a new perspective on the complicated interactions between virus and host. Research on probiotic AMGs in viruses mainly concentrates on their effects on the environment and ecosystem(23, 25–29). Gene *psbA* is considered as the biomarker for the origin classification mode of ocean water and freshwater, and can be used as a potential biomarker of classification and source tracking(25, 26). Genes *arsC* and *arsM* in soil lysogenic viruses are used to encode As (V) reduction into As (III) and to encode As (III) methylation, indicating that viral AMGs can influence bacterial metabolism through exposure to As(27). AMGs (e.g., glnK, *norB*, *nirK*, and *nirA*) carrying marine viruses potentially contribute to nitrogen cycle(23). In the low-nutrient environment in the South Pole, the viral genome contains the gene cl, which enhances hosts’ selective advantages by inhibiting expressions of pckA in the host(28, 29). These studies have revealed the advantages of AMGs in supplementing host metabolism and improving the environmental adaptation of the host. Nevertheless, there are still significant gaps concerning viruses in encoding proteins related to human metabolism or the synthesis of probiotic genes of lytic enzymes (e.g., vitamin synthase gene, glutathione synthesis gene, and cancer suppressor gene). The existence and effect of these genes on viruses have yet to be fully recognized and verified. In addition, understanding the geographic distribution patterns and functions of these probiotic genes in viruses is the basis for further understanding of the complicated interactions between viruses and hosts(30, 31).

Although some studies have revealed the functions of virus-encoded probiotic genes and virulence genes, their coexistence mechanism remain elusive(12, 13, 28, 29). Maintaining probiotic and virulence genes in viruses in different environments and host conditions and realizing their compatibility is a great challenge(32). Such a trade-off mechanism might involve complicated gene regulatory networks and may be influenced by external environmental selection stresses, such as habitat and geography(32, 33). For instance, viruses might balance the expressions of probiotic genes and virulence genes through a refined gene regulation mechanism, including transcriptional control and epigenetic modification, thus optimizing their survival and transmission abilities in different hosts and under different environmental conditions(32–34).

Based on the above review and discussion, four representative types in the literature database, namely, anticancer gene, vitamin synthesis gene, antioxidant gene, and longevity gene, were applied in this study. A total of 55 probiotic genes worth studying were searched and screened using the names of major probiotic gene types as keywords. Based on the global public database IMG/VR v4, the geological distribution and habitat features of viruses carrying probiotic gene sequences worldwide were analyzed. Moreover, viruses’ host relations and phylogenic laws were determined, and the balancing mechanism of viruses between probiotic and pathogenic functions in different environments was discussed. Furthermore, the consistency between probiotic genes in prophage gene segments of pathogenic *bacillus cereus* and probiotic AMGs of the corresponding phage was verified through single-strain whole genome and whole genome sequencing of viruses. Our results strongly prove that viruses could compensate for the functions of probiotic genes in hosts during HGT (Fig.1). We have thoroughly explored probiotic AMG sources in the virus gene library, thus this study provides new insights into the distribution, functions, and applications of viral probiotic genes.

**Fig. 1.**
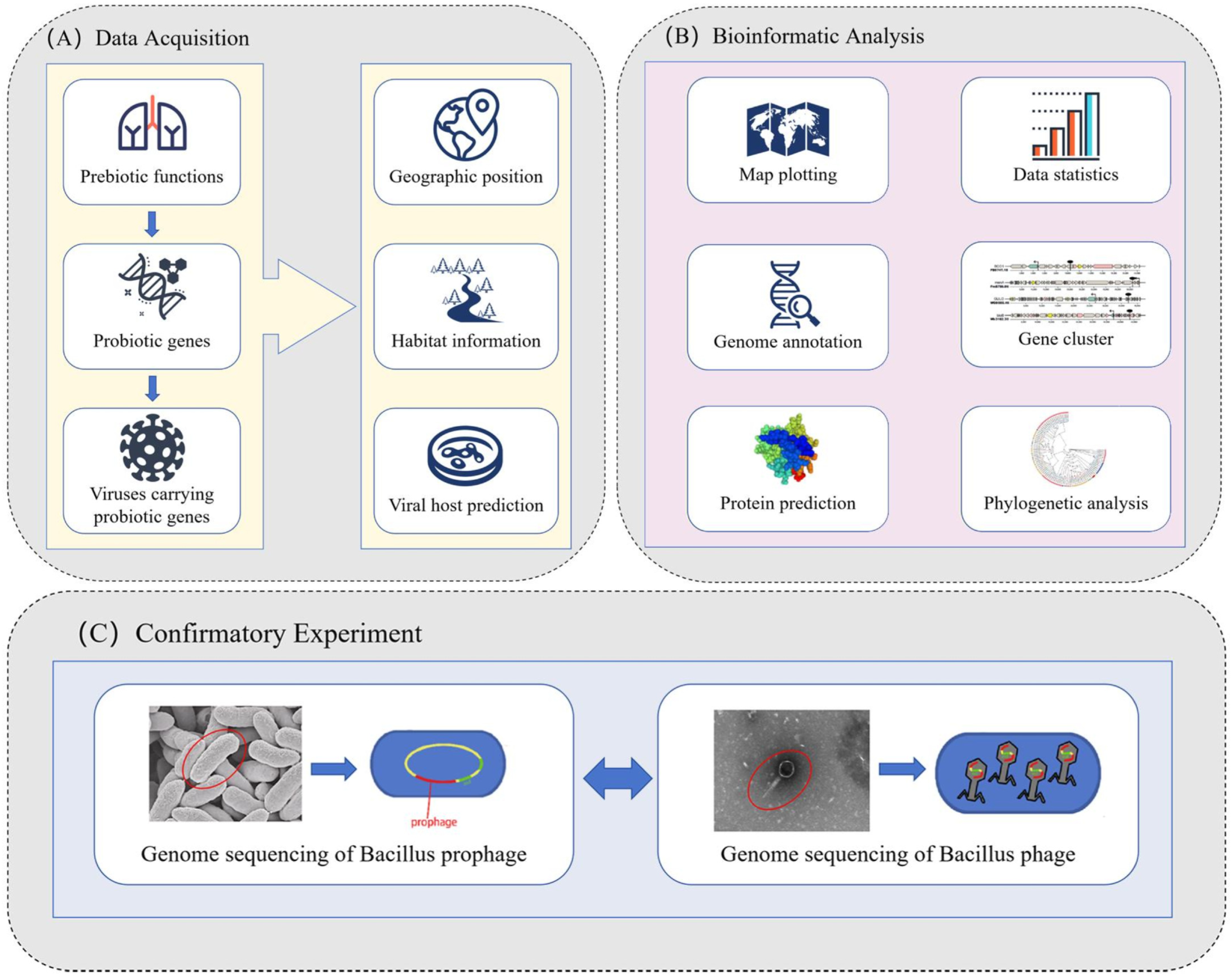
Research procedures.

## RESULTS

### Extensive distribution of viruses carrying probiotic genes

A total of 55 representative probiotic genes were screened for this study. They play an important role in the disease prevention and health continuity of humans, including anticancer, vitamin synthesis, anabolism of antioxidant substances, and longevity(35–42). Among 7,159 UVIGs, 6,583 contained geologic location information, and 6,829 contained habitat information.

This study examines viruses carrying probiotic genes distributed in 13 regions of 76 countries. Viruses that encode probiotic genes were mainly collected from America (3,866), followed by Canada (671) and the UK (258) (Fig. 2A). Viruses can be divided into eight classes: *Alsuviricetes, Caudoviricetes, Faserviricetes, Megaviricetes, Naldaviricetes, Papovaviricetes, Revtraviricetes*, and *Tectiliviricetes*. Tailed phage encoding probiotic genes accounted for the highest proportion (5,531), followed by giant viruses (1,262) (Fig. 2B). Among 40 types of probiotic genes found in virus comparison, *HSP70* (heat shock 70kDa protein) is encoded mostly through viruses (472), followed by *GPX1* (468) and *SVCT2* (448) (Fig. 2C). The sampling environments of viruses encoding probiotic genes were divided into eight classes, namely, humans, animals, freshwater, ocean water, other water bodies, soil, and plants. The highest number of viruses encoding probiotic genes (2,758) were found in freshwater, followed by the ocean (1,511) and humans (769). There were seven types of probiotic genes carrying viruses: anticancer, coenzyme Q synthesis, GSH synthesis, polyphenol synthesis, longevity, lipid-soluble vitamins, and water-soluble vitamins. Anticancer-related genes were the most common type of probiotic genes that viruses encode, amounting to 1,540. Genes related to longevity and water-soluble vitamin synthesis ranked second, amounting to 1,373 and 1,006, respectively (Fig. 2D). Research results provide theoretical support for studying the survival strategy of viruses and virus transmission paths.

**Fig. 2.**
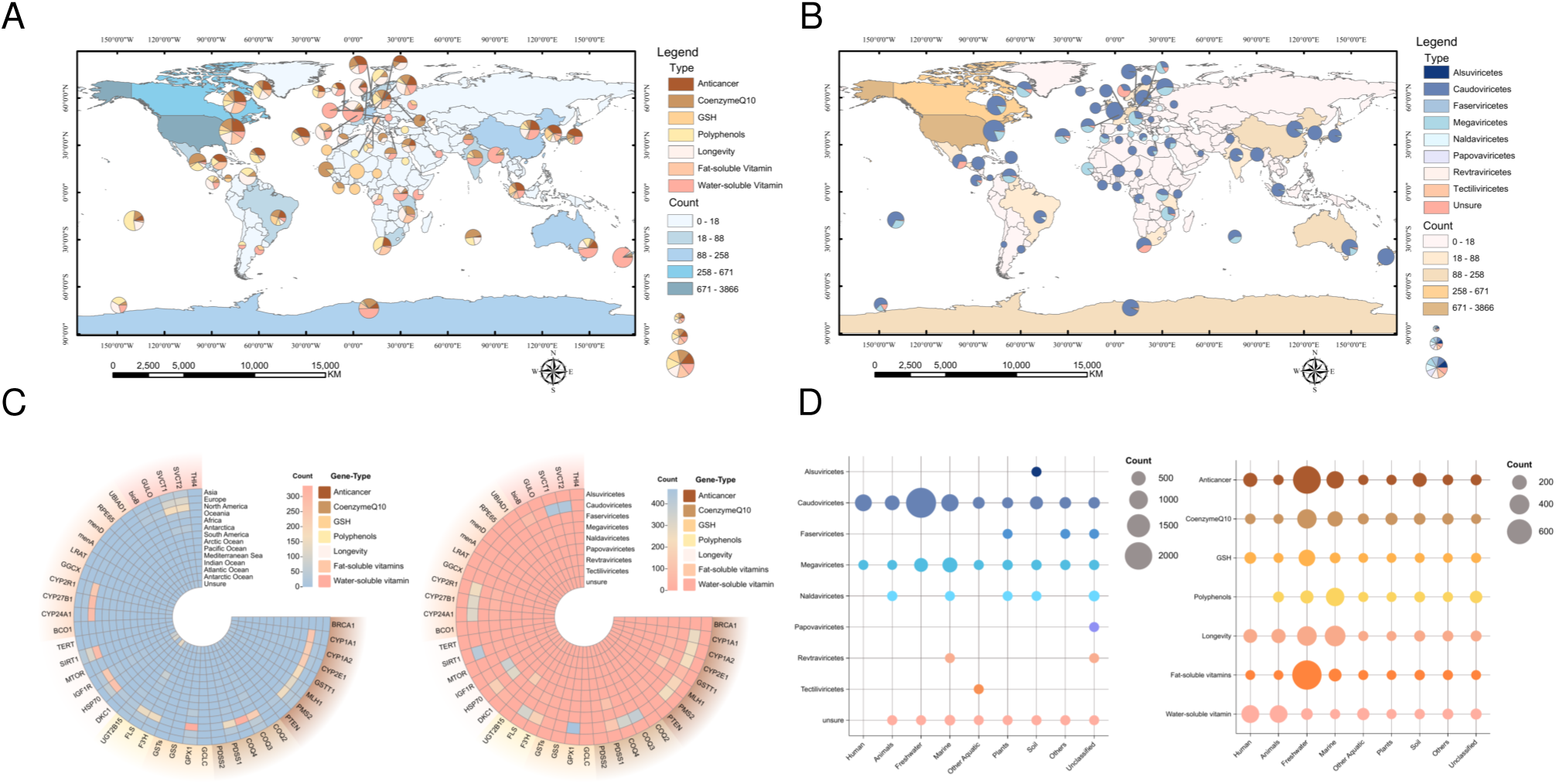
Global distribution of viruses carrying probiotic genes. A) Global distribution of viruses carry different types of probiotic genes. Different regional colors represent the abundance range of viruses carrying probiotic genes. The radius of the pie chart represents an abundance of viruses carrying probiotic genes. The pie chart represents the proportions of viruses carrying different types of probiotic genes in a region. B) Global distribution of different types of viruses includes probiotic genes. Different regional colors represent the abundance range of viruses carrying probiotic genes. The radius of the pie chart represents an abundance of viruses carrying probiotic genes. The pie chart represents the proportions of different types of viruses carrying probiotic genes in the region. C) Heat map shows the quantitative relationship between viruses carrying probiotic genes and virus classes. D) Bubble diagrams show the quantitative relationship between eight types of habitats and virus types and bubble diagrams of the quantitative relationship between habitats and gene functions.

### High host prediction rate in probiotic gene-carrying viruses in human habitats

The host prediction information of the 4,556 viral sequences in the collected data was analyzed. Among these, 904 viral sequences contain host prediction information, including 904 Kingdom and Phylum information and 896 class information (Fig. 3). Among the 904 viral sequences participating in host prediction, 26 types of hosts were predicted. Among them, 22 types belonged to the class level; four types belonged to the unpredicted class level (expressed in Phyllum), including *c_Acidimicrobiia, c_Actinomycetia, c_Alphaproteobacteria, c_Bacilli, c_Bacteroidia, c_Clostridia,* and *c_Gammaproteobacteria*. The global distribution was divided into 11 regions: Asia, Africa, Europe, North America, South America, Oceania, Antarctica, Pacific Ocean, Indian Ocean, Arctic Ocean, and Unsure. There were eight habitat distributions, including animals, freshwater, humans, marine, soil, plants, and others, which were unclassified.

**Fig. 3.**
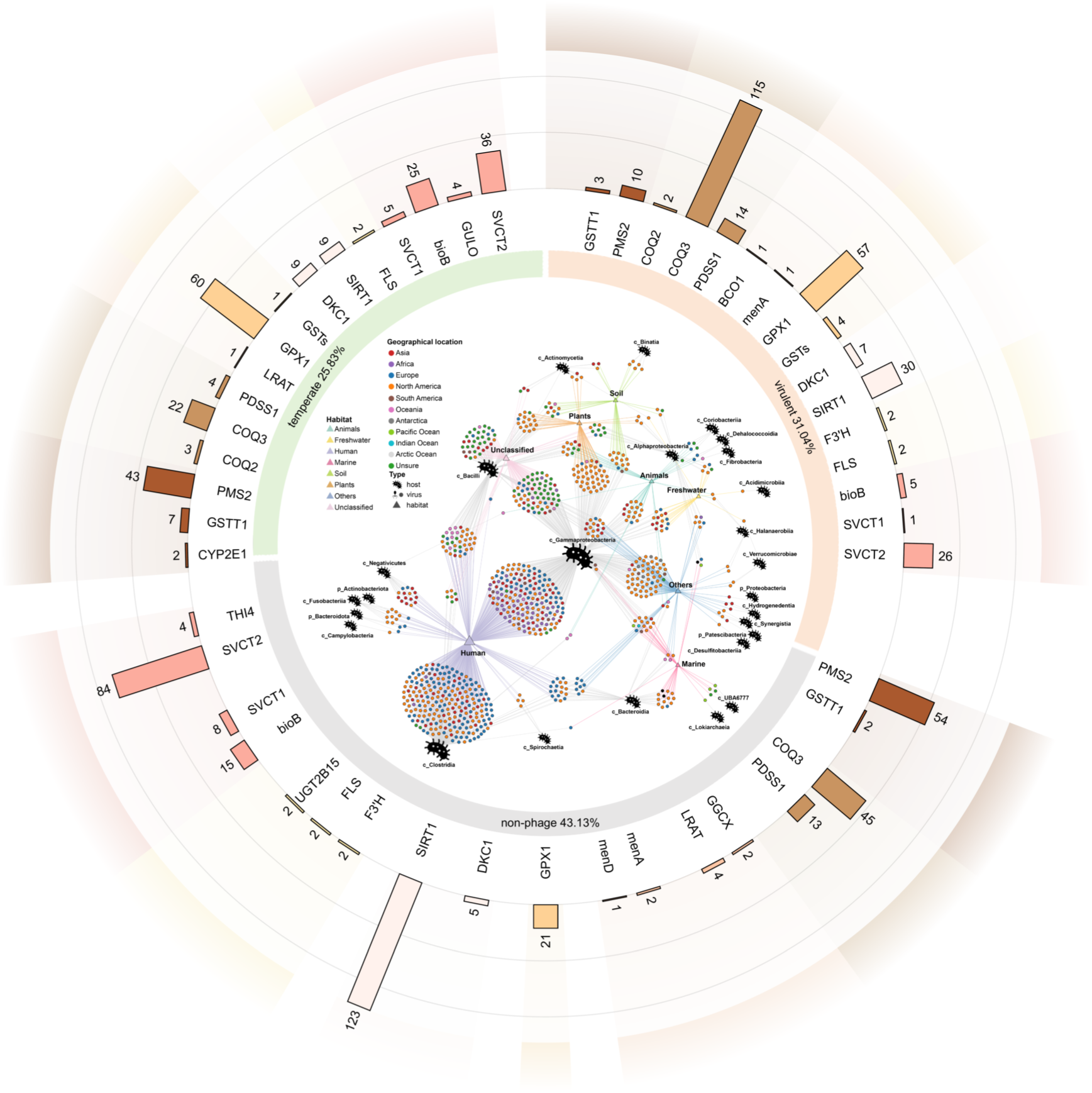
Host prediction diagram shows viruses carrying probiotic genes. Viruses were identified in equally large round charts. In the round charts, 11 colors represent global regional distributions, and the eight triangles in different colors represent the habitats. Bacterial markers represent predicted hosts. The larger marker represents the higher number of viruses belonging to this type of host or habitat. Specifically, each round point was related to the bacterial markers and triangles by two lines in different colors. The lines connecting the bacterial marker are silver, while the lines connecting with the triangles are the same color as habitats. The green of the inner ring indicates lysogenic phage, orange shows virulent phage, and gray indicates non-phages. The colors of the outer ring represent classes of probiotic genes. The histogram corresponds to the inner ring, indicating the quantity of probiotic genes in the three types of viruses.

Specifically, 389 of the 904 viral sequences were predicted as *c_Gammaproteobacteria*, accounting for the highest proportion. Next, 238 and 128 viral sequences were predicted as *c_Clostridia*, and *c_Bacilli* accounted for the second-highest proportion. Viral sequences involved in prediction were mainly collected from North America (401). This might be related to the high number of sampling sites in America. Humans are the major habitat of viral sequences involved in prediction (478), followed by plants (64). Among 514 viral sequences belonging to phage, 280 belonged to virulent phage and 233 to temperate phages. There were 321 viral sequences carrying probiotic genes in the temperate phage and 313 carrying probiotic genes in virulent phages. According to conversion, the average quantity of probiotic genes carried in each viral sequence in temperature phages is 22.3% higher than in virulent phage. The virus was divided into phages and non-phages. The phages carrying probiotic AMGs accounted for 4% of the total phages, and the non-phages carrying probiotic AMGs accounted for 1% of the total non-phages.

### Virus-encoded probiotic genes have the ability of transcription translation

Through DRAM-V and VIBRANT prediction(43, 44), this study found that *BCO1*, *bioB*, *COQ2*, *GPX1*, *GSTs*, *GSTT1*, *GULO*, and *menA* in the probiotic gene list were not only virus-encoded probiotic genes but also AMGs. A total of 133 viral sequences with high confidence levels and carrying both probiotic AMGs and virus marker genes were annotated. The phylogeny and functional expression of probiotic AMGs in viruses were analyzed. On this basis, the evolutionary relations of *GPX1*, *GSTT1*, and *bioB* in different viral sequences were disclosed (Fig. 4A). In the same region (such as North America), the pairing distance of viruses carrying the same probiotic gene *GPX1* in Marine and Freshwater ranged 0.3-0.4, showing relatively stronger consanguinity ties. When sampling sites were located in Europe, the pairing distance of viruses carrying *bioB* in humans and others was higher than 0.9, showing stronger ties of consanguinity(45–47). In particular, the pairing distance of different viruses carrying *GPX1* in humans in North America and Africa was about 0, indicating the extremely close tie of consanguinity(46–48).

**Fig. 4.**
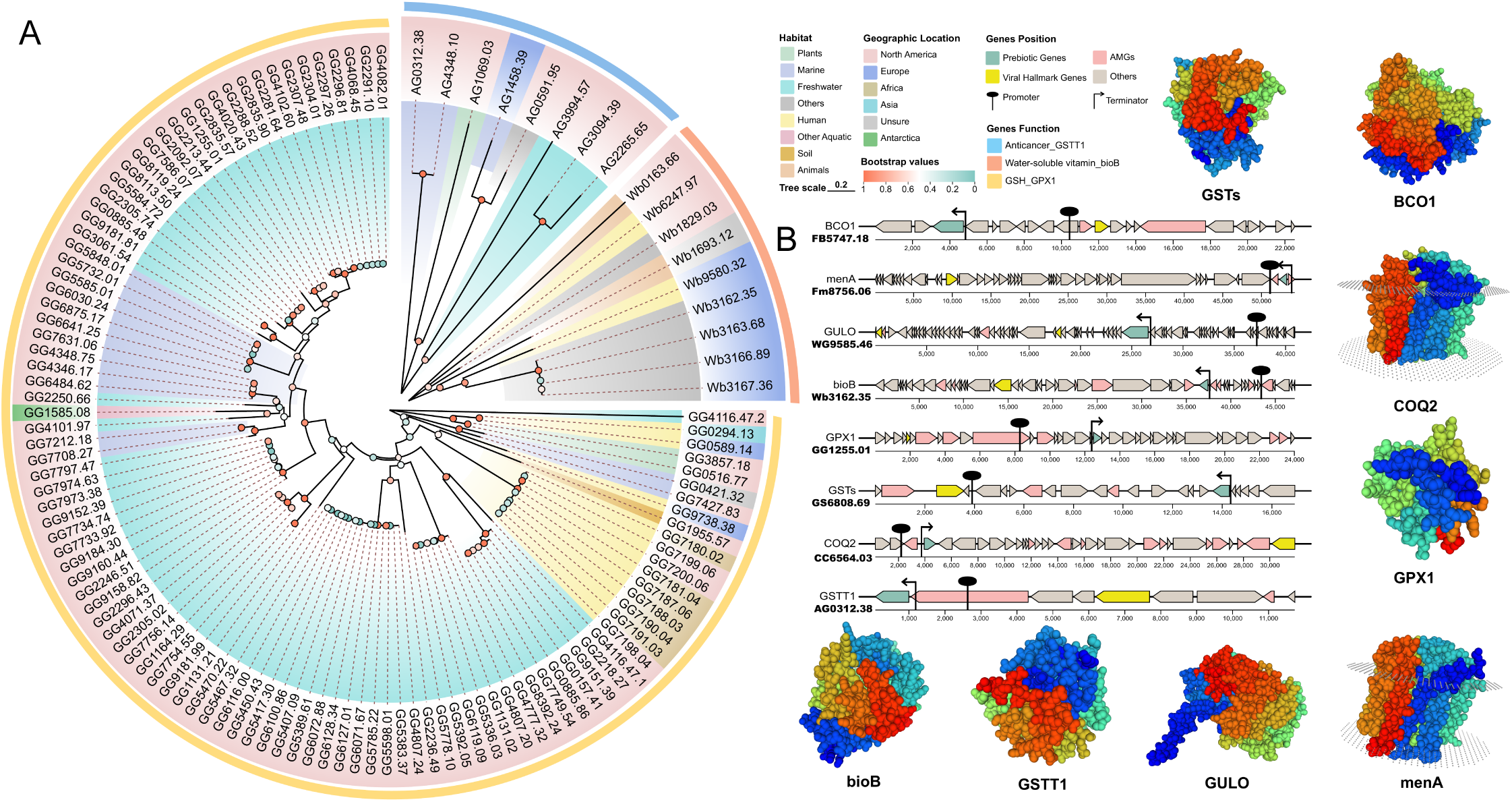
Phylogenetic tree of *GPX1*, *GSTT1*, and *bioB*, transcription translation of probiotic AMGs, and distribution of probiotic genes carried in viruses. A) Phylogenetic evolutionary relations of *GPX1*, *GSTT1*, and *bioB* are in different viral sequences. On the outer ring, *GPX1*, *GSTT1*, and *bioB* are expressed in yellow, blue, and red, respectively. B) Relative locations are shown between eight typical probiotic AMG clusters, promoters, and terminators, and four-level protein structures are predicted by probiotic AMGs. In the AMGS clusters, probiotic genes are expressed in green, virus marker genes are expressed in yellow, other AMGs are expressed in red, and non-annotated genes are expressed in gray.

In different viral sequences, promoters, and terminators who are the closest to the target could be predicted in probiotic AMGs (*BCO1, bioB, COQ2, GPX1, GSTs, GSTT1, GULO*, and *menA*), which were located in downstream of promoters and upstream of terminators. This reflects the fact that they might have transcription translation(23, 49). The structural models of these eight genes were predicted using SWISS-MODEL. The Global Model Quality Estimation (GMQE) indexes were 0.56, 0.95, 0.95, 0.80, 0.61, 0.97, 0.94, and 0.92, indicating that probiotic genes of viral sequences can encode a protein with complete structure and develop their roles (45, 50, 51) (GMQE>0.5) (Fig. 4B).

### Virulence genes in viral genomes carrying probiotic genes

In this study, four types of virulence genes (*GCH1*, *NAMPT*, *UGDH*, and *P4HA*) were recognized in viral sequences carrying probiotic genes. It was found that they could be transcripted and expressed (Fig. 5A). *GCH1* mutation can cause tetrahydrobiopterin deficiency, thus causing phenylketonuria(38, 49, 52). A high expression level of *NAMPT* might promote the growth and survival of cancer cells that participate in the development of cancer(53, 54). *UGDH* dysfunction might cause abnormal accumulation or insufficiency of the extracellular matrix, influencing tissue structures and functions. This is related to cancer and tissue fibrosis(55). The high expression *P4HA* also might be related to tumor progress and metastasis(56). Nevertheless, the GMQE values of the four types of virulence genes were 0.63, 0.53, 0.59, and 0.67 (Fig. 5B), which were generally lower than those of probiotic genes, indicating their lower possibility of transcription translation compared to probiotic genes(45, 50, 51). In this study, there was a coexistence between virulence genes and probiotic genes. According to statistics, the distance between the two genes was about 1kb–2kb (Fig. 5C).

**Fig. 5.**
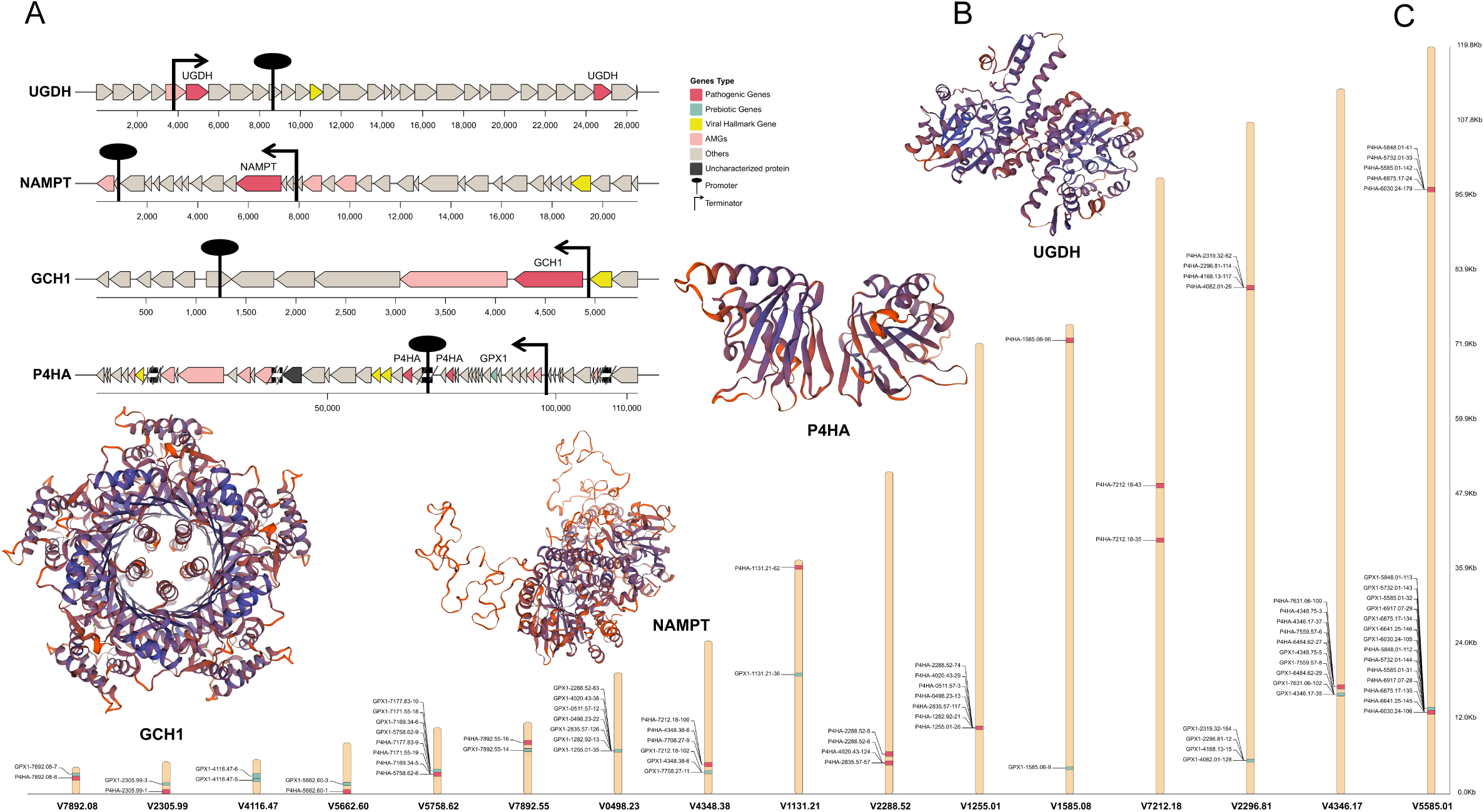
Expression potential of virulence genes and their coexistence with probiotic genes. A) Virulence gene clusters in viruses are shown carrying probiotic genes. Virulence genes are in red, probiotic genes are in green, viral marker genes are in dark red, other AMGs are in blue, non-annotated genes are in yellow, and annotated genes whose functions have not been recognized are in gray. B) Four-level protein structures are predicted by virulence genes. C) Coexistence between *P4HA* and *GPX1* is in the viral sequences.

### Consistency of probiotic genes in the whole genome sequencing of Bacillus cereus and Bacillus phages

To verify the existence and potential expression ability of probiotic genes, whole genome sequencing was carried out on phages of bacillus cereus and bacillus. It can be seen from scanning electron microscope (SEM) images that the bacillus is rod-shaped and has rough cell walls. The cell length is about 2-3μm, and the diameter is about 0.5-1μm. The SEM of the bacillus phage shows that its form includes a polyhedral head and a long tail. The diameter at the head is about 50-60nm, and the length of the tail is about 100-150nm (Fig. 6A). Phages enter host cells through the adhesion and penetration of cell walls during invasion into bacillus. Subsequently, phage genomes were integrated and entered the host genome, especially the prophage part, thus getting and expressing these probiotic genes(57–59) (Fig. 6B). According to sequencing, the prophage contained *asd* and *pdxS* in the bacillus genome, and their protein structures were predicted. The GMQE values of *asd* and *pdxS* were 0.87 and 0.86, respectively (Fig. 6C). In the genome of the bacillus phages, *asd*, and *mef* with consistent sequences were tested simultaneously (Fig. 6 D).

**Fig. 6.**
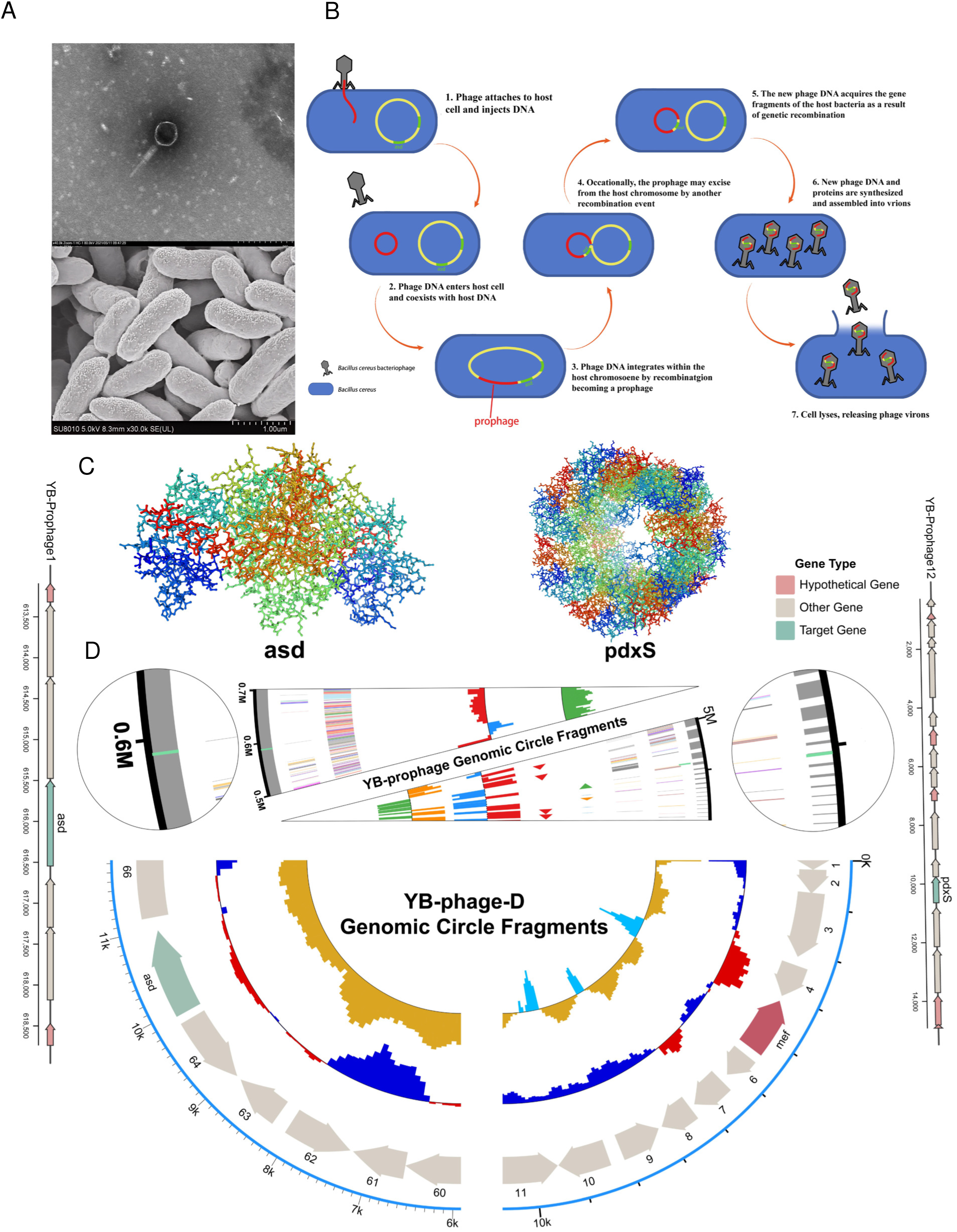
SEM and whole genome sequencing of bacillus and bacillus phages. A) SEM images of the bacillus and bacillus phages are shown in sequencing. B) The diagram shows phages invading bacillus to acquire probiotic genes. C) Sequencing disk segments of bacillus and prediction of corresponding phage gene clusters and protein structures are expressed here. D) Sequencing disk segments are shown of bacillus phages. Probiotic genes are expressed in green, virulence genes in red, and other genes in gray.

## Discussion

### Global habitat distribution and phylogenic development of probiotic genes carried in viruses

According to the global map of viral biogeography (Fig. 2), more viruses carrying probiotic gene segments in freshwater were detected than those in other habitats, accounting for 38.5% of the total viruses carrying probiotic genes. The reasons are explained as follows: The rich organic matter and microbial diversity in freshwater provide more hosts and appropriate ecological niches for these viruses, thus increasing their abundance and detection opportunities(60, 61). The number of viruses carrying probiotic gene segments collected in America is the highest, accounting for 54.0% of the total viruses. The results showed a significant preference in geological distribution and habitat selection, accompanied by diversity and universality of types and functions. In addition, it is common to have anticancer-related genes in these viruses, which might reflect the biological evolutionary strategy of viruses with advantages in host survival and adaptation in ecosystems(62, 63). Alternatively, human activities may increase the occurrence rate of cancers in the environment, thus further promoting screening and retaining of anticancer gene during the evolution of viruses to cope with such ecosystem changes(64, 65). It is important to note that although the number of viruses carrying probiotic gene segments in America is the highest, statistics indicate that the average number of probiotic genes carried in each viral sequence in Honduras was the highest (reaching 3.33), followed by that in South Africa (2.62). The number of probiotic genes in each viral sequence in America was 1.50. This might reflect the high biodiversity and complicated but stable ecosystem in Honduras, which could significantly raise the quantity of probiotic genes carried in viruses(66). In addition, the high-density population and poor public health conditions in South Africa might promote viruses to screen and retain relevant probiotic genes(67, 68). In the future, more attention will be paid to viral genome composition in regions with suitable ecological environments but low research inputs and high population density, aiming to propose and formulate scientific and objective global distribution laws of probiotic genes in viruses and provide more experimental data.

According to the data, viruses carrying the same probiotic genes in the same habitat are highly similar because they originate from common ancestral genes. Probiotic genes play a critical role in the adaptation and survival of viruses(69). As a result, these genes show high conservation in similar habitats. Given the same habitat, geological position determines the ties of consanguinity of viruses carrying probiotic genes. In North America, viruses carrying *GPX1* show close ties of consanguinity in marine and freshwater (pairing distance: 0.3-0.4). This indicates that the habitats of these water environments have small distances, and viral genome variations are highly consistent. Nevertheless, viruses carrying *bioB* in Europe show stronger ties of consanguinity in humans and other habitats (pairing distance>0.9), showing significant habitat isolation and genome differentiation. In particular, viruses carrying *GPX1* in humans in North America and Africa show almost the same genome features, indicating the extremely high similarity of viruses in these two regions at the genome level. This might be related to historical flows of people, environmental selective pressure, or similar ecological conditions(70). Therefore, this study emphasized the profound influence of geological position and habitats on the evolution of the viral genome.

### The compatible balancing mechanism between virulence genes and probiotic genes in viruses

In the confirmatory experiment, two probiotic genes (*asd* and *pdxS*) in prophages of the bacillus genome involved cell wall synthesis and synthesis of vitamin B6(71, 72). This reveals that the bacillus genome has some probiotic potentials. GMQE was predicted to be higher than 0.5, indicating that *asd* and *pdxS* have some transcription translation abilities(45, 50, 51). Moreover, the coexistence of *asd* and *mef* was detected in the genome of the bacillus phage. This reflects that phages acquire *asd* from prophages of the host bacillus through HGT. Additionally, the existence of *mef* in the phage genome reflects the potential toxicity during infection. Such gene coexistence and regulation mechanisms disclose that phages make balanced regulation between probiotic effect and pathogenic effect during long-term evolution through synchronous carrying and expressions of probiotic genes and virulence genes.

The coexistence between virulence genes and probiotic genes in viruses might originate from the cascade effect of genes(73). Thus, virulence genes and probiotic genes are influenced by similar regulation mechanisms or are acquired by viruses together through HGT(74). They are kept together in the viral genome to strengthen survival competitiveness. In the long-term evolution, viruses carefully regulate the expressions of virulent genes and probiotic genes’ balancing mechanisms to balance the probiotic and pathogenic effects. From the general perspective of the viral genome, viruses carrying virulent genes can infect and use hosts more effectively. In this way, viruses can infect the host and be duplicated, thus ensuring their transmission and survival in hosts. Virulence genes can increase the pathogenicity of viruses and help them break the hosts’ defense system, increasing transmission efficiency(75). In contrast, carrying probiotic genes can strengthen the hosts’ adaptation and survival ability, increasing the transmission potential of viruses in hosts. For example, some phages integrate probiotic genes into host genomes after infecting the host, thus increasing the environmental adaptation ability of hosts(76, 77). This is beneficial for the survival and reproduction of hosts and for the long-term transmission and survival of viruses indirectly. Therefore, viruses have realized dynamic balance in infecting hosts, using host resources and enhancing host adaptation by carrying and regulating virulence and probiotic genes. Such a balancing mechanism strengthens the survival competitiveness of viruses and discloses the complicated symbiosis and competitive relationship between viruses and hosts(78, 79).

Such a balancing mechanism reflects the adaptive adjustment of viruses during long-term evolution with hosts. If both viruses and hosts realize a dynamic balance, it can increase the transmission capacity and survival range of viruses and virus-induced diseases(80, 81). It is recommended to further study the balancing mechanism between virulence genes and probiotic genes to lower the probability of virulence gene expressions and weaken the risks of negative effects of viruses on human health. Furthermore, it can improve human health to the maximum extent by promoting the expression of probiotic genes in viruses in humans through regulatory genes, inhibiting the transcription translation of pathogenic genes in viruses, and maintaining the balanced expression of pathogenic genes and probiotic genes(82, 83).

## MATERIALS AND METHODS

### Literature review and data review

Peer review articles published in the past five decades were retrieved by databases like X-MOL (https://www.x-mol.com) and Web Of Science (https://www.webofscience.com/wos), and searching keywords included anticancer gene, vitamin synthesis gene, antioxidant gene, and longevity gene. A total of 58 studies were screened to support the study on 55 probiotic genes, mainly including 18 vitamin synthesis metabolic genes (e.g., *bioB*(35) and *menA*(36)), 14 antioxidant genes (e.g., *COQ2*(37, 38) and *GPX1*(39)), 14 anticancer genes (e.g., *GSTT1*(40)), and 9 longevity genes (e.g., *SIRT1*(41) and *IGF1R*(42)).

### Acquisition and comparison of sequences

Amino acid and nucleotide sequences of relevant genes were downloaded from the KEGG database(84). Gene names, KO number, and function information were collected. The target amino acid sequences were compared through the IMG/VR v4 database(85), obtaining the global viral sequences encoding probiotic genes (E-value < 0.0001). A total of 8,999 high-quality sequences (Bit Score > 60, Identity > 20%) were screened, finally obtaining 4,556 non-repeated viral sequences and 548 high-quality sequences(86, 87). Global spatial visualization was carried out through ArcGIS 10.7.

### Genome scanning and annotation

Viral genomes were scanned by HMMER (v3.1b2)(88). The protein sequences were compared with the HMM model of the ViPhOG database, and the Hit value with the highest scores was chosen(87, 88). The datasets were further filtered and annotated through DRAM-V (v1.2.0), thus determining the marker genes of the viruses(43). AMGs were identified by combining DRAM-V and vitality (v1.2.0)(43). The four-level structure of proteins was predicted using the SWISS-Model server (89) (https://swissmodel.expasy.org/). GMQE was higher than 0.5(45, 50, 51). Promoters and terminators were predicted through the BPROM and FindTerm server of the Softberry website(90, 91) (http://www.softberry.com/). The promoters with LDF>2.75 and terminators with the highest confidence level were screened(90, 91).

### Phylogenetic analysis

Multiple sequence comparisons of high-quality amino acid sequences were carried out using ClustalW of MEGA11(92, 93). In addition, a phylogenetic tree was built through the neighbor-joining method, and its reliability was evaluated through bootstrap analysis (1,000 repetitions) (46). The positional relationship between virulence genes and probiotic genes was exhibited by the Gene Location Visualize of TBtools(94).

### Strain cultivation and sequencing

Bacillus cereus was separated from the sewage suspension and cultivated under 37 °C, 5% CO₂. Genome DNA was extracted and quantified. Standard DNA was used to establish the library. DNA was divided into 300∼500 bp by the Covaris M220 device, followed by filling-in, adding A at the 3’ end, and connecting to the joint of TruSeq™ Nano DNA Sample Prep Kit (Turner BioSystems Inc., Sunnyvale, CA). After PCR amplification and agarose gel recovery, 2x150 bp sequencing of the DNA library was carried out on the Illumina NovaSeq 6000 platform. After quality control, sequencing data were assembled by ABySS v2.2.0, and gaps were filled with GapCloser v1.12. Moreover, encoding genes and ncRNA were annotated, and a comparative genome analysis was completed, including collinearity, SNP, and InDel.

## REFERENCES

1. Madsen A, Dai Y-N, McMahon M, Schmitz AJ, Turner JS, Tan J, Lei T, Alsoussi WB, Strohmeier S, Amor M, Mohammed BM, Mudd PA, Simon V, Cox RJ, Fremont DH, Krammer F, Ellebedy AH. 2020. Human Antibodies Targeting Influenza B Virus Neuraminidase Active Site Are Broadly Protective. Immunity 53:852–863.e7.

2. Sudhan SS, Sharma P. 2020. Chapter 4 - Human Viruses: Emergence and Evolution, p. 53–68. In Ennaji, MM (ed.), Emerging and Reemerging Viral Pathogens. Academic Press.

3. Furuse Y, Oshitani H. 2020. Viruses That Can and Cannot Coexist With Humans and the Future of SARS-CoV-2. Front Microbiol 11.

4. Long JS, Giotis ES, Moncorgé O, Frise R, Mistry B, James J, Morisson M, Iqbal M, Vignal A, Skinner MA, Barclay WS. 2016. Species difference in ANP32A underlies influenza A virus polymerase host restriction. Nature 529:101–104.

5. Delamater PL, Street EJ, Leslie TF, Yang YT, Jacobsen KH. 2019. Complexity of the Basic Reproduction Number (R0). Emerging Infectious Diseases journal.

6. Bos KI, Schuenemann VJ, Golding GB, Burbano HA, Waglechner N, Coombes BK, McPhee JB, DeWitte SN, Meyer M, Schmedes S, Wood J, Earn DJD, Herring DA, Bauer P, Poinar HN, Krause J. 2011. A draft genome of Yersinia pestis from victims of the Black Death. Nature 478:506–510.

7. Barry JM, Viboud C, Simonsen L. 2008. Cross-protection between successive waves of the 1918-1919 influenza pandemic: epidemiological evidence from US Army camps and from Britain. J Infect Dis 198:1427–1434.

8. Sharp PM, Hahn BH. 2011. Origins of HIV and the AIDS pandemic. Cold Spring Harb Perspect Med 1:a006841.

9. Lu R, Zhao X, Li J, Niu P, Yang B, Wu H, Wang W, Song H, Huang B, Zhu N, Bi Y, Ma X, Zhan F, Wang L, Hu T, Zhou H, Hu Z, Zhou W, Zhao L, Chen J, Meng Y, Wang J, Lin Y, Yuan J, Xie Z, Ma J, Liu WJ, Wang D, Xu W, Holmes EC, Gao GF, Wu G, Chen W, Shi W, Tan W. 2020. Genomic characterisation and epidemiology of 2019 novel coronavirus: implications for virus origins and receptor binding. The Lancet 395:565–574.

10. Meena M, Swapnil P, Zehra A, Aamir M, Dubey MK, Patel CB, Upadhyay RS. 2019. Chapter 11 - Virulence Factors and Their Associated Genes in Microbes, p. 181–208. In Singh, HB, Gupta, VK, Jogaiah, S (eds.), New and Future Developments in Microbial Biotechnology and Bioengineering. Elsevier, Amsterdam.

11. Keen EC. 2012. Paradigms of pathogenesis: targeting the mobile genetic elements of disease. Front Cell Infect Microbiol 2.

12. Kumar A, Prasoon P, Kumari C, Pareek V, Faiq MA, Narayan RK, Kulandhasamy M, Kant K. 2021. SARS-CoV-2-specific virulence factors in COVID-19. Journal of Medical Virology 93:1343–1350.

13. Khan N, Geiger JD. 2021. Role of Viral Protein U (Vpu) in HIV-1 Infection and Pathogenesis. 8. Viruses 13:1466.

14. Laanto E, Hoikkala V, Ravantti J, Sundberg L-R. 2017. Long-term genomic coevolution of host-parasite interaction in the natural environment. Nat Commun 8:111.

15. Jiang T, Guo C, Wang M, Wang M, You S, Liu Y, Zhang X, Liu H, Jiang Y, Shao H, Liang Y, McMinn A. 2020. Isolation and complete genome sequence of a novel cyanophage, S-B05, infecting an estuarine Synechococcus strain: insights into environmental adaptation. Arch Virol 165:1397–1407.

16. Gasper R, Schwach J, Hartmann J, Holtkamp A, Wiethaus J, Riedel N, Hofmann E, Frankenberg-Dinkel N. 2017. Distinct Features of Cyanophage-encoded T-type Phycobiliprotein Lyase ΦCpeT: THE ROLE OF AUXILIARY METABOLIC GENES*. Journal of Biological Chemistry 292:3089–3098.

17. Thompson LR, Zeng Q, Kelly L, Huang KH, Singer AU, Stubbe J, Chisholm SW. 2011. Phage auxiliary metabolic genes and the redirection of cyanobacterial host carbon metabolism. Proceedings of the National Academy of Sciences 108:E757–E764.

18. Thompson LR, Zeng Q, Kelly L, Huang KH, Singer AU, Stubbe J, Chisholm SW. 2011. Phage auxiliary metabolic genes and the redirection of cyanobacterial host carbon metabolism. Proc Natl Acad Sci U S A 108:E757–764.

19. Crummett LT, Puxty RJ, Weihe C, Marston MF, Martiny JBH. 2016. The genomic content and context of auxiliary metabolic genes in marine cyanomyoviruses. Virology 499:219–229.

20. Breitbart M, Thompson LR, Suttle CA, Sullivan MB. 2015. Exploring the Vast Diversity of Marine Viruses. Oceanography.

21. Focardi A, Ostrowski M, Goossen K, Brown MV, Paulsen I. 2020. Investigating the Diversity of Marine Bacteriophage in Contrasting Water Masses Associated with the East Australian Current (EAC) System. Viruses 12:317.

22. Lindell D, Jaffe JD, Johnson ZI, Church GM, Chisholm SW. 2005. Photosynthesis genes in marine viruses yield proteins during host infection. Nature 438:86–89.

23. Gazitúa MC, Vik DR, Roux S, Gregory AC, Bolduc B, Widner B, Mulholland MR, Hallam SJ, Ulloa O, Sullivan MB. 2021. Potential virus-mediated nitrogen cycling in oxygen-depleted oceanic waters. ISME J 15:981–998.

24. Mann NH, Cook A, Millard A, Bailey S, Clokie M. 2003. Bacterial photosynthesis genes in a virus. Nature 424:741–741.

25. Mann NH, Cook A, Millard A, Bailey S, Clokie M. 2003. Marine ecosystems: bacterial photosynthesis genes in a virus. Nature 424:741.

26. Clokie MRJ, Shan J, Bailey S, Jia Y, Krisch HM, West S, Mann NH. 2006. Transcription of a “photosynthetic” T4-type phage during infection of a marine cyanobacterium. Environ Microbiol 8:827–835.

27. Tang X, Yu P, Tang L, Zhou M, Fan C, Lu Y, Mathieu J, Xiong W, Alvarez PJ. 2019. Bacteriophages from Arsenic-Resistant Bacteria Transduced Resistance Genes, which Changed Arsenic Speciation and Increased Soil Toxicity. Environ Sci Technol Lett 6:675–680.

28. Brum JR, Hurwitz BL, Schofield O, Ducklow HW, Sullivan MB. 2016. Seasonal time bombs: dominant temperate viruses affect Southern Ocean microbial dynamics. ISME J 10:437–449.

29. Chen Y, Golding I, Sawai S, Guo L, Cox EC. 2005. Population Fitness and the Regulation of Escherichia coli Genes by Bacterial Viruses. PLOS Biology 3:e229.

30. Chakraborty C, Bhattacharya M, Sharma AR, Dhama K. 2022. Evolution, epidemiology, geographical distribution, and mutational landscape of newly emerging monkeypox virus. Geroscience 44:2895–2911.

31. Chakraborty C, Sharma AR, Bhattacharya M, Agoramoorthy G, Lee S-S. 2021. Evolution, Mode of Transmission, and Mutational Landscape of Newly Emerging SARS-CoV-2 Variants. mBio 12:10.1128/mbio.01140-21.

32. Baquero DP, Medvedeva S, Martin-Gallausiaux C, Pende N, Sartori-Rupp A, Tachon S, Pedron T, Debarbieux L, Borrel G, Gribaldo S, Krupovic M. 2024. Stable coexistence between an archaeal virus and the dominant methanogen of the human gut. Nat Commun 15:7702.

33. Sun L, You J, Li D, Zhang Z, Qin X, Pang W, Li P, Han Q, Li Y, Huang Z, Zhang X, Gong M, Yang H. 2023. Variants of a putative baseplate wedge protein extend the host range of Pseudomonas phage K8. Microbiome 11:18.

34. Albarnaz JD, Ren H, Torres AA, Shmeleva EV, Melo CA, Bannister AJ, Brember MP, Chung BY-W, Smith GL. 2022. Molecular mimicry of NF-κB by vaccinia virus protein enables selective inhibition of antiviral responses. Nat Microbiol 7:154–168.

35. Cleary PP, Campbell A, Chang R. 1972. Location of promoter and operator sites in the biotin gene cluster of Escherichia coli. Proc Natl Acad Sci U S A 69:2219–2223.

36. Miao-miao LUO, Xue-chao HU, Hong-wei QIU, Wei-ming LIU, Lu-jing REN, He H. 2015. Research progress on vitamin K2 by microorganism fermentation. 10. Food and Fermentation Industries 41:221.

37. Zhu J-Y, Fu Y, Richman A, Zhao Z, Ray PE, Han Z. 2017. A Personalized Model of COQ2 Nephropathy Rescued by the Wild-Type COQ2 Allele or Dietary Coenzyme Q10 Supplementation. J Am Soc Nephrol 28:2607–2617.

38. Mollet J, Giurgea I, Schlemmer D, Dallner G, Chretien D, Delahodde A, Bacq D, de Lonlay P, Munnich A, Rötig A. 2007. Prenyldiphosphate synthase, subunit 1 (PDSS1) and OH-benzoate polyprenyltransferase (COQ2) mutations in ubiquinone deficiency and oxidative phosphorylation disorders. J Clin Invest 117:765–772.

39. Schwarz M, Löser A, Cheng Q, Wichmann-Costaganna M, Schädel P, Werz O, Arnér ES, Kipp AP. 2023. Side-by-side comparison of recombinant human glutathione peroxidases identifies overlapping substrate specificities for soluble hydroperoxides. Redox Biol 59:102593.

40. Zhuo X, Cai L, Xiang Z, Li Q, Zhang X. 2009. GSTMI and GSTTI polymorphisms and nasopharyngeal cancer risk: an evidence-based meta-analysis. Journal of Experimental & Clinical Cancer Research 28:46.

41. Xu C, Wang L, Fozouni P, Evjen G, Chandra V, Jiang J, Lu C, Nicastri M, Bretz C, Winkler JD, Amaravadi R, Garcia BA, Adams PD, Ott M, Tong W, Johansen T, Dou Z, Berger SL. 2020. SIRT1 is downregulated by autophagy in senescence and ageing. Nat Cell Biol 22:1170–1179.

42. Abdellatif M, Trummer-Herbst V, Heberle AM, Humnig A, Pendl T, Durand S, Cerrato G, Hofer SJ, Islam M, Voglhuber J, Ramos Pittol JM, Kepp O, Hoefler G, Schmidt A, Rainer PP, Scherr D, von Lewinski D, Bisping E, McMullen JR, Diwan A, Eisenberg T, Madeo F, Thedieck K, Kroemer G, Sedej S. 2022. Fine-Tuning Cardiac Insulin-Like Growth Factor 1 Receptor Signaling to Promote Health and Longevity. Circulation 145:1853–1866.

43. Shaffer M, Borton MA, McGivern BB, Zayed AA, La Rosa SL, Solden LM, Liu P, Narrowe AB, Rodríguez-Ramos J, Bolduc B, Gazitúa MC, Daly RA, Smith GJ, Vik DR, Pope PB, Sullivan MB, Roux S, Wrighton KC. 2020. DRAM for distilling microbial metabolism to automate the curation of microbiome function. Nucleic Acids Res 48:8883–8900.

44. Kieft K, Zhou Z, Anantharaman K. 2020. VIBRANT: automated recovery, annotation and curation of microbial viruses, and evaluation of viral community function from genomic sequences. Microbiome 8:90.

45. Patil SA, Akki AJ, Raghu AV, Kulkarni RV, Akamanchi KG. 2023. Sugarcane polyphenol oxidase: Structural elucidation using molecular modeling and docking analyses. Process Biochemistry 134:243–249.

46. Saitou N, Nei M. 1987. The neighbor-joining method: a new method for reconstructing phylogenetic trees. Mol Biol Evol 4:406–425.

47. Liu L, Wu S, Yu L. 2015. Coalescent methods for estimating species trees from phylogenomic data. Journal of Systematics and Evolution 53:380–390.

48. Sourdis J, Nei M. 1988. Relative efficiencies of the maximum parsimony and distance-matrix methods in obtaining the correct phylogenetic tree. Mol Biol Evol 5:298–311.

49. Niederwieser A, Blau N, Wang M, Joller P, Atarés M, Cardesa-Garcia J. 1984. GTP cyclohydrolase I deficiency, a new enzyme defect causing hyperphenylalaninemia with neopterin, biopterin, dopamine, and serotonin deficiencies and muscular hypotonia. Eur J Pediatr 141:208–214.

50. Kiefer F, Arnold K, Künzli M, Bordoli L, Schwede T. 2009. The SWISS-MODEL Repository and associated resources. Nucleic Acids Research 37:D387–D392.

51. Liang T, Jiang C, Yuan J, Othman Y, Xie X-Q, Feng Z. 2022. Differential performance of RoseTTAFold in antibody modeling. Briefings in Bioinformatics 23:bbac152.

52. Ichinose H, Ohye T, Matsuda Y, Hori T, Blau N, Burlina A, Rouse B, Matalon R, Fujita K, Nagatsu T. 1995. Characterization of Mouse and Human GTP Cyclohydrolase I Genes: MUTATIONS IN PATIENTS WITH GTP CYCLOHYDROLASE I DEFICIENCY (∗). Journal of Biological Chemistry 270:10062–10071.

53. Kozako T, Ohsugi T, Uchida Y, Yoshimitsu M, Ishitsuka K, Kato N, Sato K, Aikawa A, Honda S. 2019. 505PHigh NAMPT expression and anti-tumour activity of NAMPT inhibitor in adult T-cell leukemia/lymphoma. Annals of Oncology 30.

54. Tang S, Garzon Sanz M, Smith O, Krämer A, Egbase D, Caton PW, Knapp S, Butterworth S. 2023. Chemistry-led investigations into the mode of action of NAMPT activators, resulting in the discovery of non-pyridyl class NAMPT activators. Acta Pharmaceutica Sinica B 13:709–721.

55. Harrington BS, Kamdar R, Ning F, Korrapati S, Caminear MW, Hernandez LF, Butcher D, Edmondson EF, Traficante N, Hendley J, Gough M, Rogers R, Lourie R, Shetty J, Tran B, Elloumi F, Abdelmaksoud A, Nag ML, Mazan-Mamczarz K, House CD, Hooper JD, Annunziata CM, Australian Ovarian Cancer Study. 2023. UGDH promotes tumor-initiating cells and a fibroinflammatory tumor microenvironment in ovarian cancer. Journal of Experimental & Clinical Cancer Research 42:270.

56. Elia I, Rossi M, Stegen S, Broekaert D, Doglioni G, van Gorsel M, Boon R, Escalona-Noguero C, Torrekens S, Verfaillie C, Verbeken E, Carmeliet G, Fendt S-M. 2019. Breast cancer cells rely on environmental pyruvate to shape the metastatic niche. Nature 568:117–121.

57. Chen J, Quiles-Puchalt N, Chiang YN, Bacigalupe R, Fillol-Salom A, Chee MSJ, Fitzgerald JR, Penadés JR. 2018. Genome hypermobility by lateral transduction. Science 362:207–212.

58. Rao VB, Feiss M. 2015. Mechanisms of DNA Packaging by Large Double-Stranded DNA Viruses. Annual Review of Virology 2:351–378.

59. Lennox ES. 1955. Transduction of linked genetic characters of the host by bacteriophage P1. Virology 1:190–206.

60. Antiqueira PAP, Petchey OL, Santos VP dos, Oliveira VM de, Romero GQ. Environmental change and predator diversity drive alpha and beta diversity in freshwater macro and microorganisms.

61. Tseng C-H, Chiang P-W, Shiah F-K, Chen Y-L, Liou J-R, Hsu T-C, Maheswararajah S, Saeed I, Halgamuge S, Tang S-L. 2013. Microbial and viral metagenomes of a subtropical freshwater reservoir subject to climatic disturbances. ISME J 7:2374–2386.

62. Rao S, Schieber AMP, O’Connor CP, Leblanc M, Michel D, Ayres JS. 2017. Pathogen-Mediated Inhibition of Anorexia Promotes Host Survival and Transmission. Cell 168:503–516.e12.

63. Correa AMS, Howard-Varona C, Coy SR, Buchan A, Sullivan MB, Weitz JS. 2021. Revisiting the rules of life for viruses of microorganisms. Nat Rev Microbiol 19:501–513.

64. Giraudeau M, Sepp T, Ujvari B, Ewald PW, Thomas F. 2018. Human activities might influence oncogenic processes in wild animal populations. Nat Ecol Evol 2:1065–1070.

65. Wu H, Eckhardt CM, Baccarelli AA. 2023. Molecular mechanisms of environmental exposures and human disease. Nat Rev Genet 24:332–344.

66. Howard AL. 1936. The Forests and Flora of British Honduras. Nature 138:145–146.

67. Achoki T, Sartorius B, Watkins D, Glenn SD, Kengne AP, Oni T, Wiysonge CS, Walker A, Adetokunboh OO, Babalola TK, Bolarinwa OA, Claassens MM, Cowden RG, Day CT, Ezekannagha O, Ginindza TG, Iwu CCD, Iwu CJ, Karangwa I, Katoto PD, Kugbey N, Kuupiel D, Mahasha PW, Mashamba-Thompson TP, Mensah GA, Ndwandwe DE, Nnaji CA, Ntsekhe M, Nyirenda TE, Odhiambo JN, Asante KO, Parry CDH, Pillay JD, Schutte AE, Seedat S, Sliwa K, Stein DJ, Tanser FC, Useh U, Zar HJ, Zühlke LJ, Mayosi BM, Hay SI, Murray CJL, Naghavi M. 2022. Health trends, inequalities and opportunities in South Africa’s provinces, 1990–2019: findings from the Global Burden of Disease 2019 Study. J Epidemiol Community Health 76:471–481.

68. de Villiers K. 2021. Bridging the health inequality gap: an examination of South Africa’s social innovation in health landscape. Infectious Diseases of Poverty 10:19.

69. Lebeer S, Vanderleyden J, De Keersmaecker SCJ. 2008. Genes and Molecules of Lactobacilli Supporting Probiotic Action. Microbiology and Molecular Biology Reviews 72:728–764.

70. Massarat A, Gymrek M, McStay B, Jónsson H. 2023. Human pangenome supports analysis of complex genomic regions. Nature 617:256–258.

71. Narayanan K, Warburton PE. 2003. DNA modification and functional delivery into human cells using Escherichia coli DH10B. Nucleic Acids Research 31:e51.

72. Leuendorf JE, Osorio S, Szewczyk A, Fernie AR, Hellmann H. 2010. Complex Assembly and Metabolic Profiling of Arabidopsis thaliana Plants Overexpressing Vitamin B6 Biosynthesis Proteins. Molecular Plant 3:890–903.

73. Fotsing SF, Margoliash J, Wang C, Saini S, Yanicky R, Shleizer-Burko S, Goren A, Gymrek M. 2019. The impact of short tandem repeat variation on gene expression. Nat Genet 51:1652–1659.

74. Anderson RP, Roth JR. 1978. Tandem chromosomal duplications in *Salmonella typhimurium*: Fusion of histidine genes to novel promoters. Journal of Molecular Biology 119:147–166.

75. Geoghegan JL, Holmes EC. 2018. The phylogenomics of evolving virus virulence. Nat Rev Genet 19:756–769.

76. Chevallereau A, Pons BJ, van Houte S, Westra ER. 2022. Interactions between bacterial and phage communities in natural environments. Nat Rev Microbiol 20:49–62.

77. Feiner R, Argov T, Rabinovich L, Sigal N, Borovok I, Herskovits AA. 2015. A new perspective on lysogeny: Prophages as active regulatory switches of bacteria. Nature reviews Microbiology 13:641–650.

78. Duggal NK, Emerman M. 2012. Evolutionary conflicts between viruses and restriction factors shape immunity. Nat Rev Immunol 12:687–695.

79. Roossinck MJ. 2011. The good viruses: viral mutualistic symbioses. Nat Rev Microbiol 9:99–108.

80. Jarett JK, Džunková M, Schulz F, Roux S, Paez-Espino D, Eloe-Fadrosh E, Jungbluth SP, Ivanova N, Spear JR, Carr SA, Trivedi CB, Corsetti FA, Johnson HA, Becraft E, Kyrpides N, Stepanauskas R, Woyke T. 2020. Insights into the dynamics between viruses and their hosts in a hot spring microbial mat. ISME J 14:2527–2541.

81. Ghazal P, García-Ramírez J, González-Armas JC, Kurz S, Angulo A. 2000. Principles of homeostasis in governing virus activation and latency. Immunol Res 21:219–223.

82. Gan L, Xu W-H, Xiong Y, Lv Z, Zheng J, Zhang Y, Lin J, Liu J, Chen S, Chen M, Guo Q, Wu J, Chen J, Su Z, Sun J, He Y, Liu C, Wang W, Verstraete W, Sorgeloos P, Defoirdt T, Qin Q, Liu Y. 2021. Probiotics: their action against pathogens can be turned around. Sci Rep 11:13247.

83. Singh K, Rao A. 2021. Probiotics: A potential immunomodulator in COVID-19 infection management. Nutrition Research 87:1–12.

84. Kanehisa M, Goto S. 2000. KEGG: Kyoto Encyclopedia of Genes and Genomes. Nucleic Acids Research 28:27–30.

85. Camargo AP, Nayfach S, Chen I-MA, Palaniappan K, Ratner A, Chu K, Ritter SJ, Reddy TBK, Mukherjee S, Schulz F, Call L, Neches RY, Woyke T, Ivanova NN, Eloe-Fadrosh EA, Kyrpides NC, Roux S. 2023. IMG/VR v4: an expanded database of uncultivated virus genomes within a framework of extensive functional, taxonomic, and ecological metadata. Nucleic Acids Research 51:D733–D743.

86. Roux S, Adriaenssens EM, Dutilh BE, Koonin EV, Kropinski AM, Krupovic M, Kuhn JH, Lavigne R, Brister JR, Varsani A, Amid C, Aziz RK, Bordenstein SR, Bork P, Breitbart M, Cochrane GR, Daly RA, Desnues C, Duhaime MB, Emerson JB, Enault F, Fuhrman JA, Hingamp P, Hugenholtz P, Hurwitz BL, Ivanova NN, Labonté JM, Lee K-B, Malmstrom RR, Martinez-Garcia M, Mizrachi IK, Ogata H, Páez-Espino D, Petit M-A, Putonti C, Rattei T, Reyes A, Rodriguez-Valera F, Rosario K, Schriml L, Schulz F, Steward GF, Sullivan MB, Sunagawa S, Suttle CA, Temperton B, Tringe SG, Thurber RV, Webster NS, Whiteson KL, Wilhelm SW, Wommack KE, Woyke T, Wrighton KC, Yilmaz P, Yoshida T, Young MJ, Yutin N, Allen LZ, Kyrpides NC, Eloe-Fadrosh EA. 2019. Minimum Information about an Uncultivated Virus Genome (MIUViG). Nat Biotechnol 37:29–37.

87. Roux S, Paul BG, Bagby SC, Nayfach S, Allen MA, Attwood G, Cavicchioli R, Chistoserdova L, Gruninger RJ, Hallam SJ, Hernandez ME, Hess M, Liu W-T, McAllister TA, O’Malley MA, Peng X, Rich VI, Saleska SR, Eloe-Fadrosh EA. 2021. Ecology and molecular targets of hypermutation in the global microbiome. Nat Commun 12:3076.

88. Eddy SR. 2011. Accelerated Profile HMM Searches. PLOS Computational Biology 7:e1002195.

89. Waterhouse A, Bertoni M, Bienert S, Studer G, Tauriello G, Gumienny R, Heer FT, de Beer TAP, Rempfer C, Bordoli L, Lepore R, Schwede T. 2018. SWISS-MODEL: homology modelling of protein structures and complexes. Nucleic Acids Research 46:W296–W303.

90. Deltcheva E, Chylinski K, Sharma CM, Gonzales K, Chao Y, Pirzada ZA, Eckert MR, Vogel J, Charpentier E. 2011. CRISPR RNA maturation by trans-encoded small RNA and host factor RNase III. Nature 471:602–607.

91. Rinaldi AJ, Lund PE, Blanco MR, Walter NG. 2016. The Shine-Dalgarno sequence of riboswitch-regulated single mRNAs shows ligand-dependent accessibility bursts. Nat Commun 7:8976.

92. Thompson JD, Gibson TobyJ, Higgins DG. 2003. Multiple Sequence Alignment Using ClustalW and ClustalX. Current Protocols in Bioinformatics 00:2.3.1–2.3.22.

93. Tamura K, Stecher G, Kumar S. 2021. MEGA11: Molecular Evolutionary Genetics Analysis Version 11. Molecular Biology and Evolution 38:3022–3027.

94. Chen C, Chen H, Zhang Y, Thomas HR, Frank MH, He Y, Xia R. 2020. TBtools: An Integrative Toolkit Developed for Interactive Analyses of Big Biological Data. Molecular Plant 13:1194–1202.

